# Colitis is associated with loss of LHPP and up-regulation of histidine phosphorylation in intestinal epithelial cells

**DOI:** 10.1101/2020.10.11.334334

**Authors:** Markus Linder, Dritan Liko, Venkatesh Kancherla, Salvatore Piscuoglio, Michael N. Hall

## Abstract

Protein histidine phosphorylation (pHis) is a posttranslational modification involved in cell cycle regulation, ion channel activity and phagocytosis (1). Using novel monoclonal antibodies to detect pHis (2), we recently reported that loss of the histidine phosphatase LHPP results in elevated pHis levels in hepatocellular carcinoma (3). Here, we show that intestinal inflammation correlates with loss of LHPP, in DSS-treated mice and in inflammatory bowel disease (IBD) patients. Increased histidine phosphorylation was observed in intestinal epithelial cells (IECs), as determined by pHis immunofluorescence staining of colon samples from a colitis mouse model. However, ablation of *Lhpp* did not cause increased pHis or promote intestinal inflammation in physiological conditions or after DSS treatment. Our observations suggest that increased histidine phosphorylation plays a role in colitis, but loss of LHPP is not sufficient to increase pHis or to cause inflammation in the intestine.

## Introduction

Protein histidine phosphorylation, a poorly characterized posttranslational modification, refers to the the addition of a phosphate group to the imidazole ring of histidine via a heat and acid labile phosphoramidate (P-N) bond. Both nitrogens in the histidine imidazole ring can be phosphorylated resulting in the formation of two isomers: 1-phosphohistidine (1-pHis) and 3-phosphohistidine (3-pHis). So far, three mammalian histidine phosphatases (LHPP, PGAM5 and PHPT1) and two histidine kinases (NME1, NME2) have been described (1). We recently reported that murine and human hepatocellular carcinomas (HCCs) exhibit elevated histidine phosphorylation and decreased levels of the histidine phosphatase LHPP. Re-introduction of LHPP resulted in decreased pHis levels in vitro and prevented tumor formation in an HCC mouse model, suggesting that elevated pHis is pathological (3). Another recent study showed that LHPP protein expression correlates with survival of colorectal cancer (CRC) patients, again suggesting that LHPP acts as a tumor suppressor (4).

Inflammatory bowel disease (IBD) is a major risk factor for CRC (5). IBD is a general term for intestinal disorders characterized by chronic colitis, such as Crohn’s disease (CD) and ulcerative colitis (UC). The etiology and the molecular pathophysiology of IBDs are incompletely understood, resulting in insufficient progress in the development of novel therapies (6). Importantly, inhibition or deletion of the K^+^ channel KCa3.1 prevents colitis progression (7), and PGAM5 inhibits KCa3.1 by dephosphorylating NME2-pHis118 (8). Taken together, the above suggests that pHis may play a role in the pathology of IBD.

## Results and Discussion

To investigate if histidine phosphorylation plays a role in colitis, we analyzed a publicly available transcriptomic profile (E-GEOD-16879) from colon samples of healthy humans and treatment-naïve IBD patients (9). Patients suffering from CD or UC showed significantly decreased expression of *LHPP*, but no difference in expression of the other known histidine phosphatase genes *PGAM5* and *PHPT1* (Fig. 1A). Expression of *NME1* and *NME2* was significantly upregulated in patients suffering from IBDs (Fig. 1A). Based on these findings, we hypothesized that elevated pHis, via down-regulation of the histidine phosphatase LHPP and up-regulation of the histidine kinases NME1/2, contributes to disease progression.

**Figure 1.**
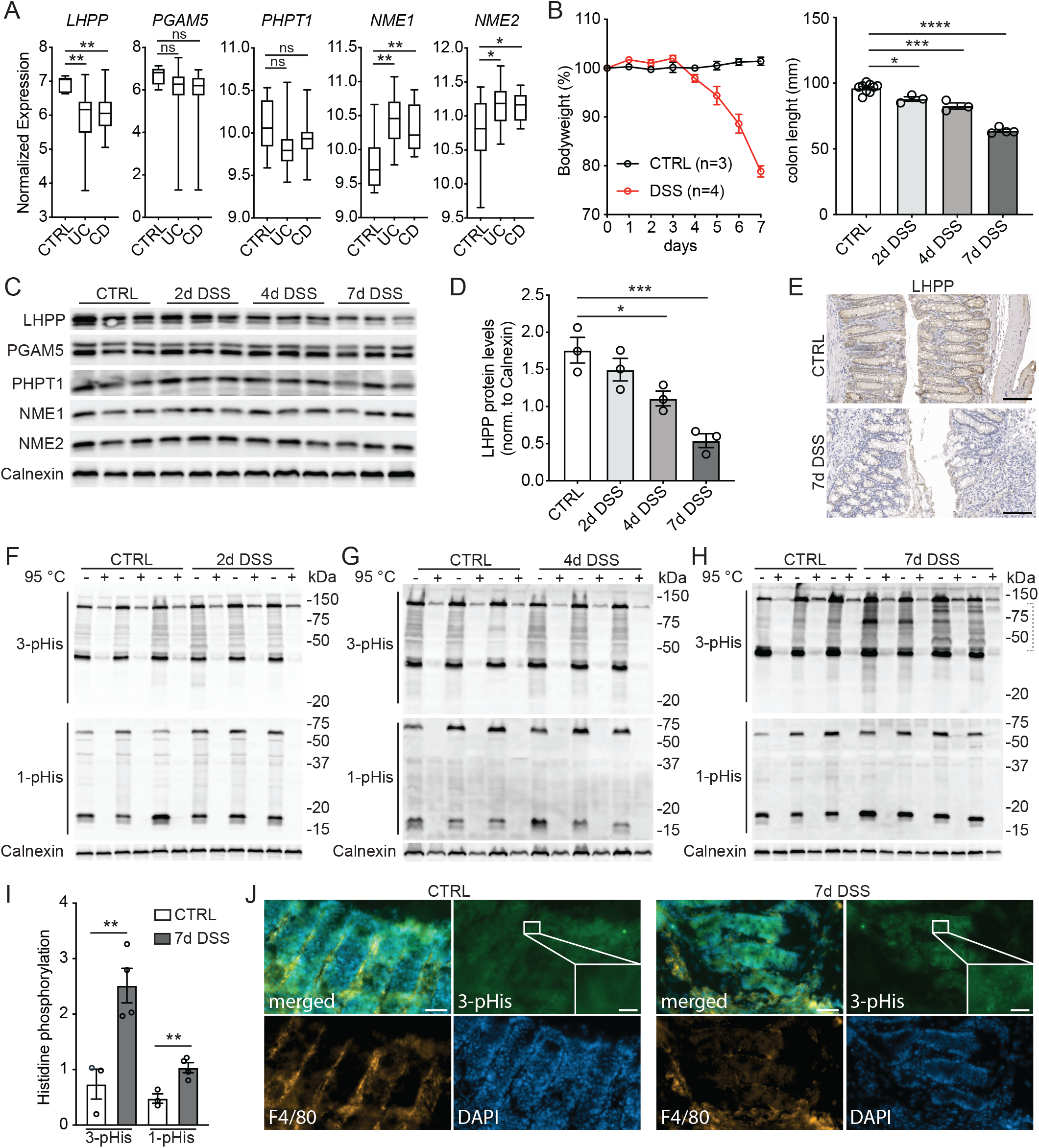
Intestinal inflammation correlates with down-regulation of LHPP and increased histidine phosphorylation. (*A*) Analysis of publicly available dataset comparing mRNA expression of known histidine phosphatases (*LHPP, PGAM5, PHPT1*) and histidine kinases (*NME1/2*) in colon tissue from healthy patients (CTRL) with patients suffering from ulcerative colitis (UC) and Crohn’s disease (CD). (*B*) Bodyweight and colon length of mice treated (DSS) and untreated (CTRL) with DSS at indicated timepoints (days). (*C*) Immunoblot analysis of histidine phosphatases and kinases in colon lysates of DSS-treated and -untreated mice at indicated timepoints. (*D*) Quantification of LHPP protein levels normalized to Calnexin at different timepoints of DSS treatment. (*E*) Immunohistochemistry visualization of LHPP in colon samples of mice treated (DSS) and untreated (CTRL) for 7 days with DSS. Scale bar: 100μm. (*F-H*) Immunoblot analysis of 1- and 3-pHis levels in colon samples from DSS-treated (DSS) and -untreated (CTRL) mice at indicated timepoints. (*I*) Quantification of 1- and 3-pHis levels in *H*. (*J*) DAPI and Immunofluorescence staining of colon samples from untreated (CTRL) and 7-day, DSS-treated mice. Scale bars: 50μm (low magnification) and 10μm (high magnification).

To examine further a role of pHis in IBD, we induced experimental colitis by treating wild-type mice with dextran sodium sulfate (DSS) (10). Mice exhibited mild colitis-like symptoms, after 2 to 4 days, that developed into severe colitis with 20% bodyweight loss and significantly decreased colon length, as observed after one week of treatment (Fig. 1B). Next, we analyzed expression of known histidine phosphatases and kinases at different timepoints of DSS treatment. The histidine phosphatase LHPP, but no other histidine phosphatase, was significantly down-regulated in the colon of wild-type mice after 4 days of DSS treatment, as determined by immunoblotting (Fig. 1C, D). After one week of treatment, LHPP expression was further reduced, indicating that LHPP expression negatively correlated with colitis severity (Fig. 1C, D). Importantly, immunohistochemistry (IHC) revealed that DSS-dependent down-regulation of LHPP was due mainly to reduced expression in IECs rather than in infiltrating immune cells (Fig. 1E). Expression of the histidine kinases was unchanged in DSS-treated mice (Fig. 1C).

We next analyzed histidine phosphorylation in colon samples isolated at different timepoints of DSS treatment. Immunoblot analysis of colon lysates showed significantly up-regulated 1- and 3-pHis levels after 7 days of DSS treatment, but not at earlier timepoints (Fig. 1F-I). In agreement with the IHC results described above, DSS treatment increased pHis exclusively in IECs, not in infiltrating macrophages (F4/80-positive cells), as shown by immunofluorescence staining (Fig. 1J). As we observed down-regulation of LHPP at 4 days of DSS treatment but changes in pHis only after 7 days, i.e., LHPP loss preceded increased pHis, we speculate that loss of LHPP contributes to high levels of pHis and disease progression.

To obtain insight on the role of LHPP in normal development and in colitis progression, we generated full body LHPP knockout (*Lhpp*^−/−^) mice using CRSPR/Cas9 (see Materials and Methods). *Lhpp*^−/−^ mice were vital, fertile and indistinguishable from littermate controls. Histological analysis of the colonic mucosa did not reveal significant differences between *Lhpp*^−/−^ and *Lhpp*^+/+^ littermates (Fig. 2A). Colons of 1.5-year-old *Lhpp*^−/−^ mice displayed no sign of cancer and/or inflammation, and no changes in proliferation or apoptosis (Fig. 2A). Moreover, immunoblot analysis revealed no significant difference in 1- or 3-pHis levels in large intestine lysates of *Lhpp*^−/−^ and *Lhpp*^+/+^ littermates (Fig. 2B, C). Colon from *Lhpp*^−/−^ mice did not display any differences in PHPT1, NME1 and NME2 protein levels (Fig. 2B, D), but did exhibit elevated PGAM5 expression, possible as a compensatory mechanism for loss of LHPP (Fig. 2B).

**Figure 2.**
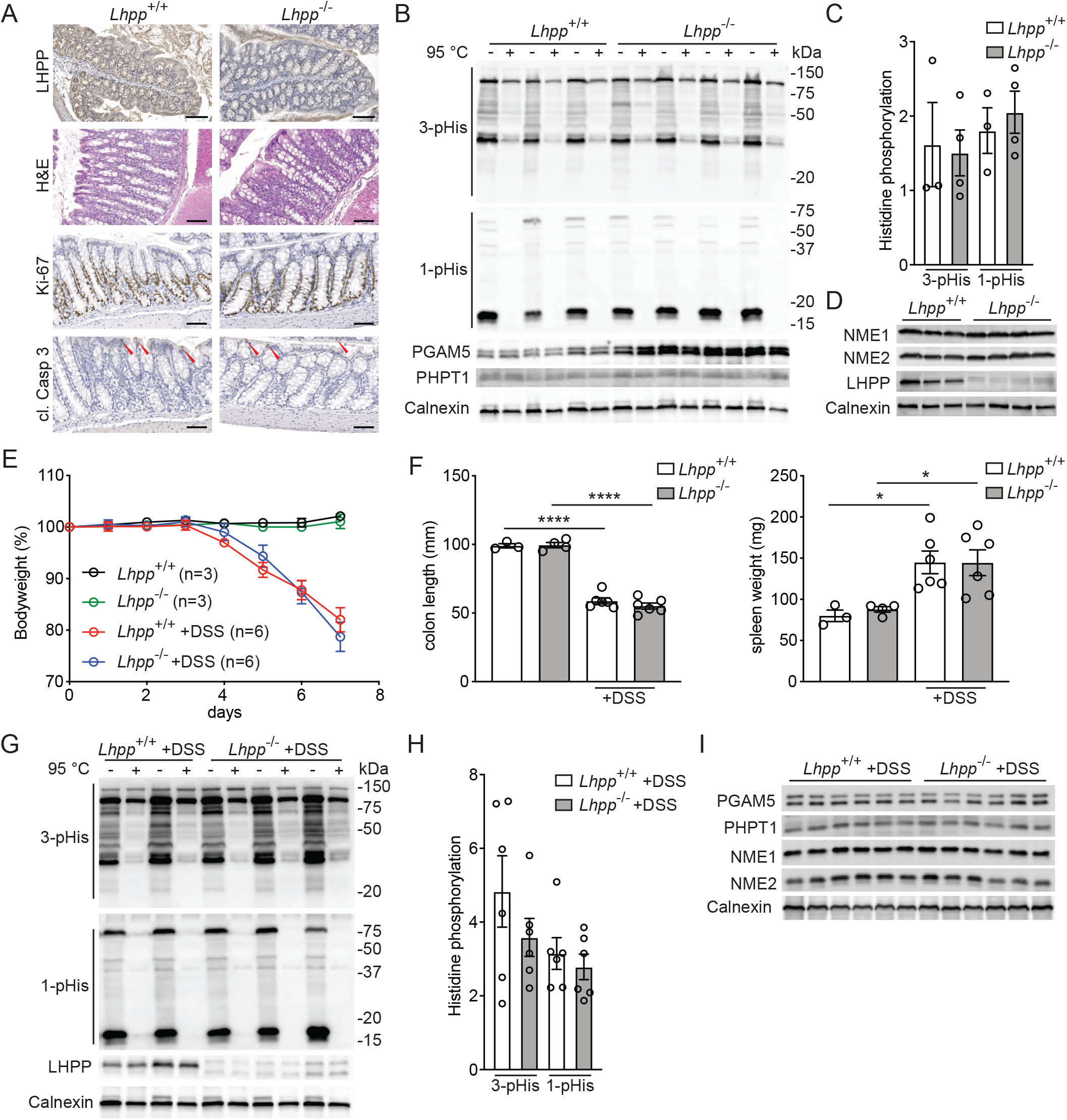
LHPP is dispensable for colitis development. (*A*) IHC and H&E stainings of colon samples of 1.5-year-old *Lhpp*^+/+^ and *Lhpp*^−/−^ mice. Red arrowheads indicate cleaved (cl.) caspase 3 positive cells. Scale bars: 100μm. (*B*) Immunoblot analysis of 1- and 3-pHis and histidine phosphatase protein levels in colon samples from *Lhpp*^+/+^ and *Lhpp*^−/−^ mice. (*C*) Quantification of 1- and 3-pHis levels in *B*. (*D*) Immunoblot analysis of NME1, NME2 and LHPP in colon samples from *Lhpp*^+/+^ and *Lhpp*^−/−^ mice. (*E*) Bodyweight during DSS treatment. (*F*) Colon length and spleen weight after 7 days of DSS treatment. (*G*) Immunoblot analysis of 1- and 3-pHis and histidine phosphatase protein levels in colon samples from *Lhpp*^+/+^ and *Lhpp*^−/−^ mice treated with DSS for 7 days. (H) Quantification of 1- and 3-pHis levels in *G* with n=6. (*I*) Immunoblot analysis of histidine phosphatases and kinases in colon samples *Lhpp*^+/+^ and *Lhpp*^−/−^ mice treated with DSS for 7 days.

To determine whether loss of LHPP has an impact on colitis progression, we treated *Lhpp*^−/−^ and *Lhpp*^+/+^ littermates with DSS. All colitis-related parameters such as weight loss, colon shrinkage, and elevated spleen weight were similar in *Lhpp*^−/−^ and *Lhpp*^+/+^ mice, indicating that LHPP loss does not impact colitis development (Figs. 2E, F). Finally, we did not detect genotype-dependent changes in intestinal 1- and 3-pHis levels after DSS treatment (Fig. 2G, H), or differences in expression of known histidine phosphatases or kinases (Fig. 2I).

In summary, we show that LHPP loss and increased histidine phosphorylation in intestinal cells correlate with colitis. However, LHPP loss does not appear to be sufficient for either the observed increase in pHis or inflammation, at least in mice. Increased NME1/2 expression or activity, which we observed in IBD patients but not in *Lhpp*^−/−^ mice, might be required in addition to loss of LHPP. Little is known about the role of pHis in inflammation. Fuhs et al reported that both malignant epithelial cells and macrophages show high pHis levels in vitro, and suggested that pHis is important in phagocytosis (2). As activated macrophages are key players in colitis (11), we originally expected that the high pHis levels we observed in our experimental system would be in immune cells. However, results from our IF staining indicate that DSS treatment triggers pHis in IECs. As we observed high pHis levels only in late-stage colitis, we cannot exclude that IECs up-regulate pHis as a response to infiltrating immune cells. Alternatively, upregulated pHis levels in epithelial cells might promote inflammation by inducing the production of pro-inflammatory signaling molecules and recruitment of macrophages. It remains to be determined whether increased pHis is a consequence or a cause of inflammation in IBD. It is also necessary to investigate the role of pHis in the complex interplay between IECs and immune cells. Finally, it will be important to identify histidine phosphorylated proteins and to determine their role in inflammatory diseases.

## Materials and Methods

### Mice

To generate *Lhpp*^−/−^ mice, exon 2 of the mouse *Lhpp* gene was deleted using CRISPR/Cas9-mediated non-homologous end joining (NHEJ). Two gRNAs were designed to target the *Lhpp* introns 1 (IVS1) and 2 (IVS2). The sequences targeting the respective introns IVS1-catctgactcacatcatgtgagg and IVS2-gcatcctgaagctagccttgagg were selected for optimal on target activity using the CRISPOR online tool (12). NHEJ events at the gRNA target sites led to the excision of the genomic fragment containing exon 2 resulting in a *Lhpp*-null allele. CRISPR/Cas9-mediated modification of the *Lhpp* sequence was carried out by electroporation of fertilized mouse oocytes as previously described (13). *Lhpp*^−/−^ and *Lhpp*^+/+^ mice were maintained in a C57BL/6J genetic background. C57BL/6J wild-type mice were purchased from Janvier Labs. Experimental colitis was induced in 8− to 12-week-old male mice by administering 2.5% DSS in drinking water for up to 7 days according to published protocols (10). All animal experiments conducted were compliant with federal laws and guidelines and were approved by the veterinary office of Basel-Stadt.

### Histology

Immunohistochemistry (IHC) and H&E stainings were performed as previously described (14). The following primary antibodies were used: cleaved Caspase 3 (9664; CST), Ki-67 (12202; CST), LHPP (NBP1-83272; Novus). For Immunofluorescence (IF) stainings, colons were flushed with ice cold PBS (pH 8.5), cryo-fixed in OCT and stored at −80°C. To prevent heat- and acid-mediated pHis degradation, all steps during the IF staining process were performed at 4°C and the pH of all buffers was adjusted to 8.5. After cutting, colon cryo-sections (10μm) were fixed for 1h in 4% PFA, washed with 1x PBS, blocked for 1h using blocking buffer (1x PBS, 1% BSA, 0.05% Triton-X 100) and subsequently incubated O/N with primary antibodies (3-pHis: rabbit, SC44-1 and F4/80: rat, ab6640; abcam) diluted in blocking buffer. Afterwards, slides were rinsed with PBS and incubated for 1h with secondary antibodies (Alexa Fluor 488 anti-rabbit and Alexa Fluor 568 anti-rat; Invitrogen) and DAPI (4083; CST). Finally, the stained sections were washed and mounted with water-based mounting medium (H-1400; Vector Laboratories).

### Immunoblotting

Immunoblots were performed and quantified as previously described (3). To detect pHis, the following monoclonal primary antibodies were used: 1-pHis (0.5ug/ml, SC1-1), 3-pHis (0.5ug/ml, SC44-1). For regular immunoblot analysis the following primary antibodies were used: Calnexin (ADI-SPA-860; Enzo), LHPP (15759-1-AP; Proteintech), NME1 (3345; CST), NME2 (ab60602; abcam) PGAM5 (ab126534; abcam), PHPT1 (LS⍰C192376; LSBio).

### Analysis of publicly available transcriptomic dataset

For mRNA expression analysis the Affymetrix GeneChip Human Genome U133 Plus 2.0 gene expression datasets E-GSE16879 (9) was downloaded from ArrayExpress. The probes were matched with gene names, using biomaRt R package. Afterwards, the gene expression levels of *LHPP*, *PGAM5*, *PHPT1 NME1* and *NME2* were analyzed between different conditions (CTRL, UC, CD).

### Statistical analysis

Data analysis was performed with PRISM 8.0 (GraphPad). Single comparisons were performed by unpaired, 2-tailed students t-test. Comparison of multiple groups were performed by one-way ANOVA followed by Tukey’s post hoc test for multiple comparison. Data are shown as mean ◻±◻SEM. \**P* < 0.05, \*\**P* < 0.01, \*\*\**P* < 0.001, \*\*\*\**P* < 0.001.

## Author Contributions

M.L. and M.N.H. conceived and designed research; M.L., V.K., D.L. and S.P. performed research; M.N.H contributed resources and secured funding; M.L., V.K., D.L., S.P. and M.N.H. analyzed data; and M.L. and M.N.H. wrote the paper.

## Acknowledgments

This work was supported by an EMBO long-term fellowship to M.L., and by grants from the Swiss National Science Foundation and the European Research Council (MERiC) to M.N.H.

